# CilioGenics: an integrated method and database for predicting novel ciliary genes

**DOI:** 10.1101/2023.03.31.535034

**Authors:** Mustafa S. Pir, Ferhan Yenisert, Aslı Karaman, Efe Begar, Sofia Tsiropoulou, Elif Nur Firat-Karalar, Oliver E Blacque, Sukru S. Oner, Osman Doluca, Sebiha Cevik, Oktay I. Kaplan

## Abstract

Discovering the entire list of human ciliary genes would help in the diagnosis of cilia-related human disorders known as ciliopathy, but at present the genetic diagnosis of many ciliopathies (over 30%) is far from complete (Bachmann-Gagescu et al., 2015; Knopp et al., 2015; Paff et al., 2018). In a theory, many independent approaches may uncover the whole list of ciliary genes, but 30% of the genes on the ciliary gene list are still ciliary candidate genes (van Dam et al., 2019; Vasquez et al., 2021). All of these cutting-edge techniques, however, have relied on a different single strategy to discover ciliary candidate genes. Because different methodologies demonstrated distinct capabilities with varying quality, categorizing the ciliary candidate genes in the ciliary gene list without further evidence has been difficult. Here, we present a method for predicting ciliary capacity of each human gene that incorporates diverse methodologies (single-cell RNA sequencing, protein-protein interactions (PPIs), comparative genomics, transcription factor (TF)-network analysis, and text mining). By integrating multiple approaches, we reveal previously undiscovered ciliary genes. Our method, CilioGenics, outperforms other approaches that are dependent on a single method. Our top 500 gene list contains 256 new candidate ciliary genes, with 31 experimentally validated. Our work suggests that combining several techniques can give useful evidence for predicting the ciliary capability of all human genes.

## Introduction

Cilia are cellular organelles that protrude from the surfaces of most cells and play critical roles in cellular motility, sensation, and embryo development (Satir and Christensen, 2007). Based on the organization of their microtubule-based axoneme core (presence or absence of a central pair of microtubules) and their motility behavior, cilia are subdivided into two types: motile and non-motile (primary cilium). Different types of cilia perform distinct functions. Mucus clearance, for example, requires the coordinated beating of multiple motile cilia on epithelial cells in respiratory organs, whereas the non-motile cilium is needed for chemical, thermal, and mechanical sensations (Mitchison and Valente, 2017). Functional and structural abnormalities of cilia cause a variety of diseases, including Bardet-Biedl Syndrome (BBS), Joubert syndrome (JS), and Nephronophthisis (NPHP), known as ciliopathies, which are accompanied by a variety of symptoms such as cystic kidneys, retinal degeneration, and retinitis pigmentosa, obesity, and intellectual disability as a result of defects in multiple organs and tissues (Reiter and Leroux, 2017). Many signaling pathways, including Hedgehog, Wnt, Notch, Hippo, and PDGF, essential for cell/tissue formation and homeostasis, have been linked to cilia, and some of their components are enriched in cilia (Wheway et al., 2018).

Because of the significance of cilia for human health, many researchers have been trying to identify the parts list of this tiny and complex organelle. A variety of approaches have been used to determine the protein compositions of cilia, leading to the identification of over 600 ciliary genes and many candidate ciliary genes. These approaches include genomic comparisons of ciliary and non-ciliary organisms, the presence or absence of ciliary gene-specific transcription factor (TF) binding sites such as the X-box motif, an increase of gene expression during cilia assembly, functional genomics (for example, RNAi and CRISPR screening), clinical studies, co-expression of ciliary genes within specific tissues, and proteomics, (Andersen et al., 2003; Arnaiz et al., 2010; Avidor-Reiss et al., 2004; Baron et al., 2007; Blacque et al., 2005; Boesger et al., 2009; Boldt et al., 2016; Breslow et al., 2018; Broadhead et al., 2006; Cao et al., 2017; Choksi et al., 2014; Efimenko et al., 2005; Fritz-Laylin and Cande, 2010; Geremek et al., 2014, 2011; Hodges et al., 2011; Hoh et al., 2012; Ishikawa et al., 2012; Ivliev et al., 2012; Jakobsen et al., 2011; Keller et al., 2005; Kilburn et al., 2007; Kim et al., 2010; Kubo et al., 2008; Lambacher et al., 2016; Lauwaet et al., 2011; Li et al., 2004; Liu et al., 2007; May et al., 2021; Mayer et al., 2009, 2007; McClintock et al., 2008; Merchant et al., 2007; Mick et al., 2015; Müller et al., 2010; Nakachi et al., 2011; Nogales-Cadenas et al., 2009; Ostrowski et al., 2002; Pazour et al., 2005; Phirke et al., 2011; Reyfman et al., 2019; Roosing et al., 2015; Ross et al., 2007; Sigg et al., 2017; Stubbs et al., 2008; UK10K Consortium et al., 2015; van Dam et al., 2019; Vasquez et al., 2021; Yano et al., 2013). The constant discovery of new ciliary genes implies that these independent methods are inadequate to uncover all ciliary genes, even though they have assisted in the discovery of many ciliary genes. This is especially true for comparative genomics, which reveals new ciliary genes on the basis of being present in organisms with cilia but not organisms that lack cilia. This method, however, misses genes such as AP1M1 (Adaptor Related Protein Complex 1 Subunit Mu 1), which is found in almost all organisms’ genomes and is required for cilia biogenesis in *C. elegans* and correct cilia orientation in humans (Avidor-Reiss et al., 2004; Kaplan et al., 2010; Li et al., 2014; Nevers et al., 2017).

Given that cilia are highly specialized organelles whose functions are required in many cell types but not all cell types in humans or other organisms, cilia-enriched genes are more likely to be expressed in ciliated cells rather than non-ciliated cells. Single-cell RNA sequencing (scRNA-seq) based method, which would reveal gene expression differences among cell types, would overlook genes expressed in all types but required for cilia assembly and/or function. The main problem is that all of these independent approaches have produced far too many ciliary candidate genes, and the real question is whether there is an ideal method for discovering new ciliary genes that outperform others.

Here, we develop an integrated method, called CilioGenics by taking into account several independent approaches, including comparative genomics, scRNA-seq analysis, gene regulatory network, protein-protein interactions (PPI), and text mining to obtain a comprehensive list of potential ciliary genes. The integrated CilioGenics method is superior over any single method, allowing us to confidently identify novel ciliary genes. We confirm the ciliary localization of many new genes, including ZC2HC1A, ZNF474, WDR54, TMEM145, and TTC39C. We also made available all data to the CilioGenics website (https://ciliogenics.com/) for scientists to examine the data.

## Results

### Single cell RNA sequencing analysis of *C. elegans* reveals novel ciliary candidate genes

We integrate multiple datasets to create a novel method to accurately predict ciliary genes (**Figure 1**). *C. elegans* has a variety of cell types, including the intestine, muscle, germ cells, and neurons (cholinergic neurons and ciliated sensory neurons). Ciliary genes are exclusively expressed in ciliated sensory neurons, but not in non-ciliary cells such as intestine, muscle, germ cells, and non-ciliated neurons (Cao et al., 2017). We hypothesize that genes that are specifically expressed in ciliated sensory neurons but not in other tissues should be considered ciliary genes. Thus, to find new ciliary genes, we analyzed the single-cell RNA sequencing (scRNA-seq) data of *C. elegans* using the previously published *C. elegans* scRNA-seq data (Cao et al., 2017).

**Figure 1.**
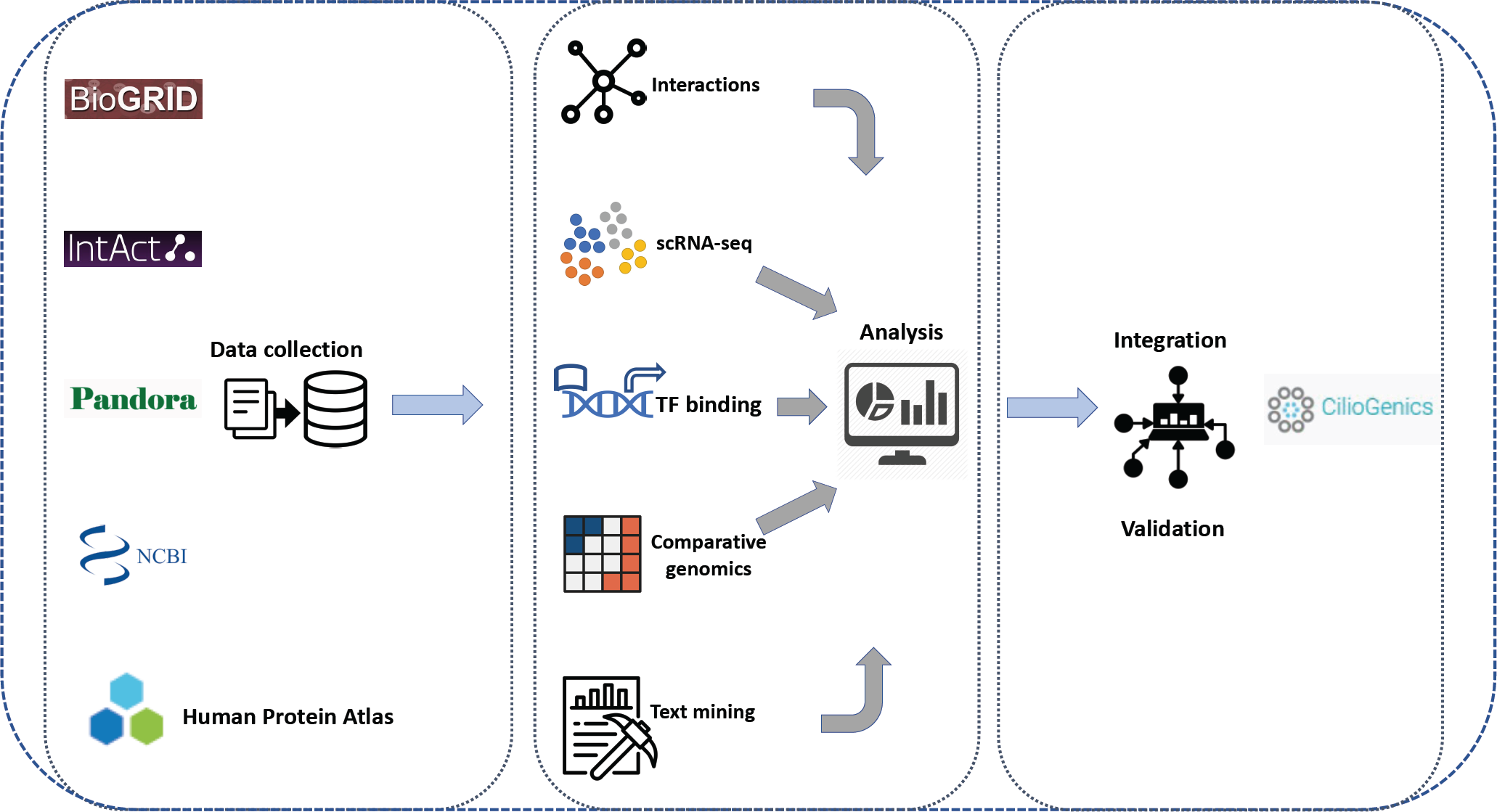
Workflow of CilioGenics. Data from IntAct, BioGRID and HuRI, Pandora, and NCBI is gathered, assessed using R scripts, and then integrated with data from other sources.

Analysis reveals that 1989 *C. elegans* genes appear to be differentially expressed and specifically enriched in ciliated sensory neurons, including amphid, phasmid, and oxygen sensory neurons, and may be good ciliary candidate genes; however, only 685 have human counterparts and those without are ignored. We discover that 379 of the 685 genes are already known ciliary genes, while the rest are classified as putative candidate ciliary genes that we are further investigating (**Table S1**). Expectedly, many core ciliary genes, including intraflagellar transport (IFT) components, transition zone (TZ), and BBSome components, exhibit exclusive ciliary cell-specific expressions (**Figure 2A**), and many previously unknown ciliary genes recapitulate the cilia-specific expression patterns of known ciliary genes. For example, our study discovers *WDR-31* (WD Repeat Domain 31) (*T05A8.5*) and ELMD-1 (ELMOD1-3) as putative ciliary genes, and we confirm the exclusive expression of both genes in the ciliated sensory neurons in *C. elegans*, as well as their localization to cilia/basal body in human cell lines and *C. elegans,* implying that our approach could reveal novel ciliary genes (Cevik et al., 2021) (**Figure 2C**). To confirm the ciliary expression of several putative genes among the ciliary gene candidates, we investigated the expressions of *tmem-145* (human *TMEM145* and *C. elegans C15A7.2*), *wdr-54* (human *WDR54* and *C. elegans* F39H12.2), and *zchc-1a* (human *ZC2HC1A* and *C. elegans T03G11.3*) in *C. elegans,* and find that they are exclusively expressed in the ciliated sensory neurons in *C. elegans.* Thus, analysis of scRNA-seq data in *C. elegans* is an excellent approach to identifying new ciliary genes.

**Figure 2.**
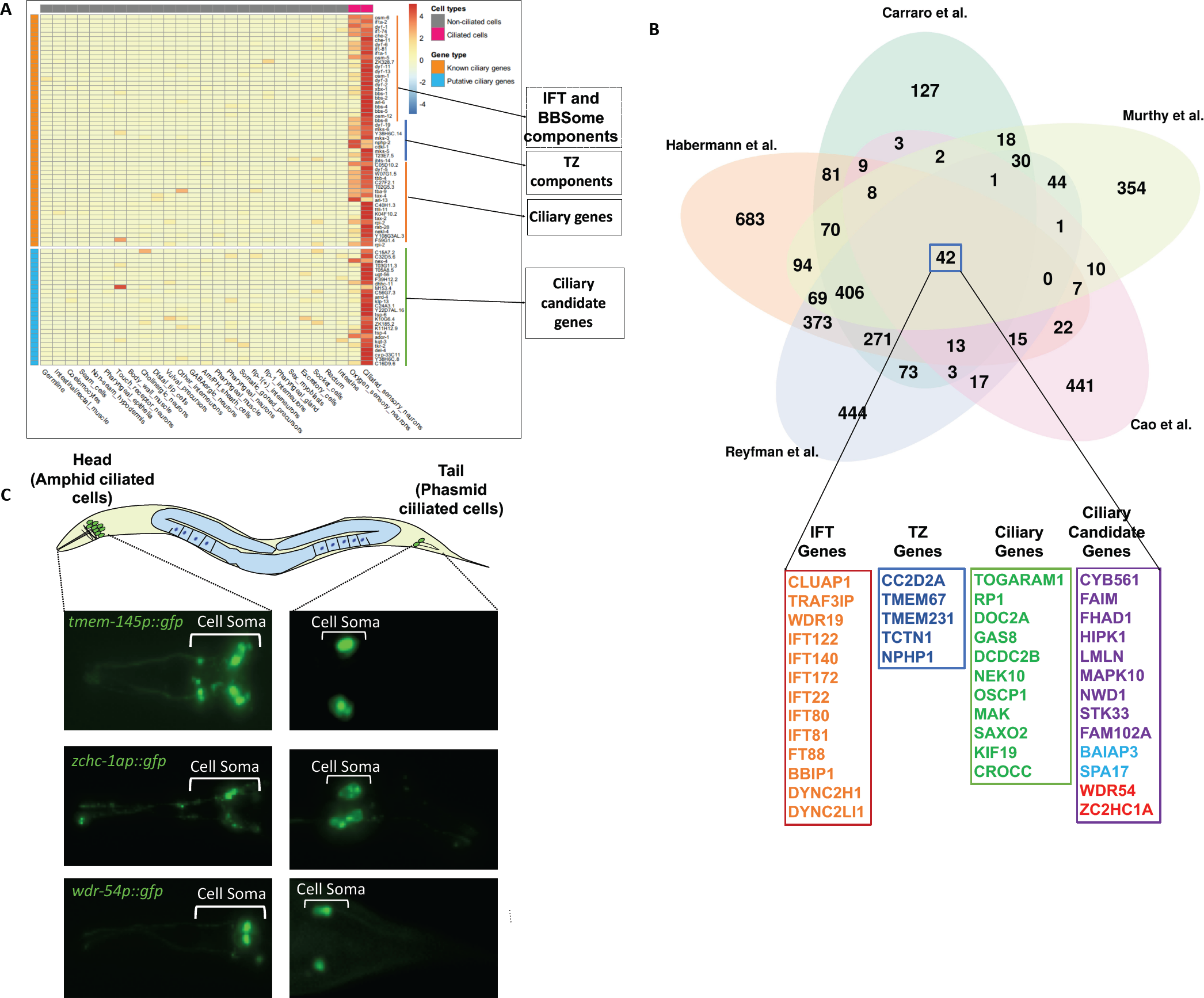
Single-cell (sc) RNA-seq of *C. elegans* reveals many ciliary candidate genes. **A)** The differentially expressed genes in the *C. elegans* scRNA-seq dataset are shown in a representative heatmap. Cells with cilia are denoted by the color red, whereas cells without cilia are denoted by the color gray. Candidate genes for cilia are shown in blue, whereas known ciliary genes are shown in orange. 42 genes shared by five studies were displayed beneath the Venn diagram. Intraflagellar transport (IFT) genes (orange), transition zone (TZ) genes (blue), ciliary genes (green), and putative ciliary genes (light blue, red, and purple) are shown. Putative ciliary gene encoding shown in light blue or red is found to localize to cilia by us or others (Ivliev et al., 2012; McClintock et al., 2008). Please see **Figure 8B** for WDR54 and ZC2HCA1 localization. **B)** Venn diagram shows the number of shared genes between the Reyfman (human lung scRNA-seq), Carraro (human lung scRNA-seq), Habermann (human lung scRNA-seq), Murthy (human lung scRNA-seq), and Cao (*C. elegans* scRNA-seq). **C)** Shown are drawings of *C. elegans*. The ciliated sensory neurons (green) in the head and tail are depicted. Head and tail fluorescence images show the expression of *wdr-54promoter::gfp, tmem-145promoter::gfp*, and *zchc-1apromoter::gfp.* Cell somas (body) of sensory neurons are depicted in white brackets.

### Analysis of four human lung scRNA-seq uncovers new ciliary candidate genes

We next analyzed scRNA-seq data from human lung tissue. Because s of its wide range of ciliary and non-ciliary cell types, including multiciliated epithelial cells, club cells, and alveolar type I (AT1) and type II (AT2) cells, the lung is an ideal system for comparing the expression profiles of cells with and without cilia (**Figure S1**). We selected four different lung scRNA-seq data sets and examined them individually and comparatively (Carraro et al., 2021; Habermann et al., 2020; Kadur Lakshminarasimha Murthy et al., 2022; Reyfman et al., 2019). An analysis of the Reyfman, Carraro, Habermann, and Murthy data sets reveals a total of 1802, 1157, 2165, and 966 genes that are expressed exclusively in ciliated cells, respectively, with 1082, 719, 1017, and 406 emerging as putative candidate ciliary genes (Reyfman: 240 known ciliary genes, Carraro: 231 known ciliary genes, Habermann: 287 known ciliary genes, Murthy: 190 known ciliary genes) (**Figure 3A, B and C, and Table S2**). Many known ciliary genes display cilia-specific expression patterns and enrichment (visit https://www.ciliogenics.com/), and three, including IFT88 (an IFT gene), TMEM231 (transition zone gene) and NEK10 (NIMA Related Kinase 10) are displayed in the UMAP plots (**Figure 3D, E, and F**). Thus, analysis of human lung scRNA-seq data identifies known and candidate ciliary genes.

**Figure 3.**
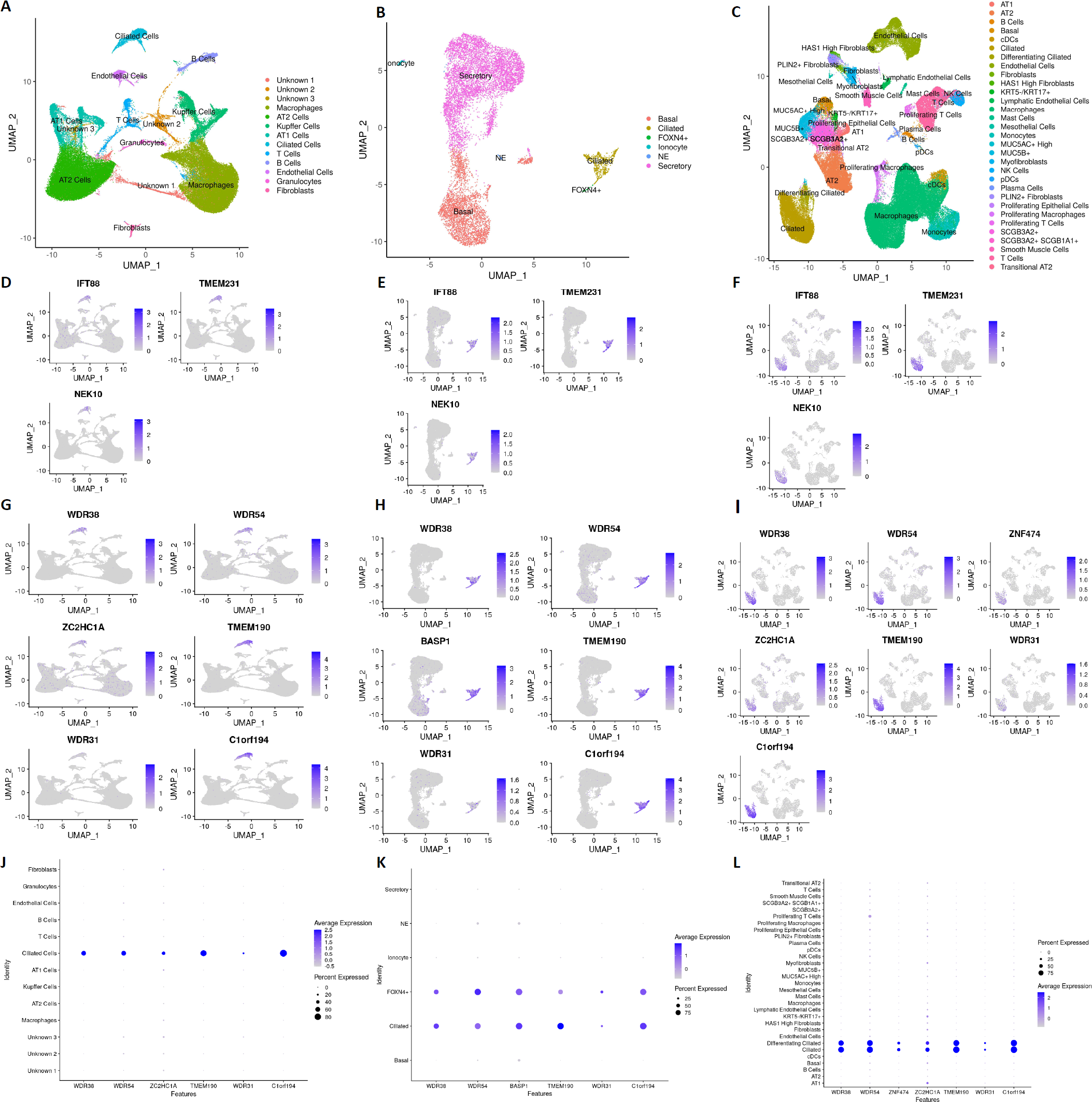
Human lung scRNA-seq reveals many ciliary candidate genes. **A, B, and A) C)** The UMAPs from Reyfman, Carraro, and Habermann’s human lung scRNA-Seq are shown in (A), (B), and (C), respectively. **D, E, and F)** UMAPs of IFT88, TMEM231, and NEK10 from scRNA-seq Reyfman, Carraro, and Habermann are presented in (D), (E), and (F), respectively. **G, H, I, J, K, and L**) UMAPs and Dotplots of the indicated ciliary candidate genes for scRNA-seq Reyfman (G and J), Carraro (H and K), and Habermann are displayed (I and L).

Comparison of the *C. elegans* scRNA-seq and the four human lung scRNA-seq ciliogenics datasets (Reyfman, Murthy, Carraro, and Habermann) reveals 42 genes in common (**Figure 2B**). This 42 gene list contains 29 known ciliary genes that function with processes such as IFT and transition zone gating, as well as 13 candidate ciliary genes. These candidates include *WDR54* and *ZC2HC1A*, which we show using GFP reporters are expressed in *C. elegans* ciliated cells (**Figure 2C**). Interestingly, for the *BAIAP3* and SPA17 candidate ciliary genes, independent studies revealed that they encode proteins that localize to cilia.

We also compared the four human scRNA-seqs datasets (Carraro, Reyfman, Habermann, and Murthy) without the *C. elegans* scRNA-seq dataset because many human genes have no orthologs in *C. elegans*. This analysis uncovers 524 genes in common, including *WDR38, TMEM190*, and *C1orf194*, which we deem as strong candidate ciliary genes. All ciliary candidate genes, including *WDR38, TMEM190,* and *C1orf194* display cilia-specific expression patterns in dot plots of all human scRNA-seqs (**Figure 3J, K and L, and** visit https://www.ciliogenics.com/ for the expression patterns of every human gene). All ciliary candidate genes discovered by scRNA-seq are included in **Table S2**. Of note, although *WDR31* does not appear a strong ciliary candidate gene based on the results of these human scRNA-seq analyses, the dot plot showed that its expression is limited to ciliated cells, although the expression level is low (**Figure 3G, H, and I**). Together with *WDR31* being highly represented in the *C. elegans* scRNA-seq dataset, and previously confirmed cilia localizations for *WDR31* (Cevik et al., 2021), it is clear that the gene is cilia-related, and further confirms our approach here to reveal novel ciliary genes. It is worth mentioning that users may access *C. elegans* and human scRNA-seq data and visualize the expression patterns of each gene on the CilioGenics website (https://ciliogenics.com/).

### Comparative genomics (phylogenetic profiling) reveals previously unidentified ciliary candidate genes

Because not all eukaryotic species need the specialized function of cilia, cilia have been lost in many organisms, including plants and fungi. The assumption is that genes with a unique role for cilia should be present in genomes of ciliated organisms but not in the genomes of organisms without cilia. For this reason, comparative genomics has been extensively employed to uncover ciliary genes. We performed comparative genomics on 72 eukaryotic species for over 20,000 human protein-coding genes to predict the human genes involved in ciliary functions and then compared the candidate ciliary genes to the candidate ciliary genes from our scRNA-seq analyses. Our comparative genomics followed by the dissimilarity matrix hierarchical clustering reveals a total of 40 gene clusters, including two cilia-specific clusters (31 and 37), average conservation clusters like vertebrate-specific clusters (1, 4, 6, 9, 16, and 30) and non-cilia-specific clusters (low specificity clusters: 2, 3, 5, 7, 8, 10, 11, 12, 13, 14, 15, 17, 18, 19, 20, 21, 22, 23, 24, 25, 26, 27, 28, 29, 31, 32, 33, 34, 35, 36, 37, 38, 39, and 40) (**Figure 4A, B, C, and D**). The average conservation clusters are believed to have potential ciliary genes, but their ciliary potential is lower than that of two cilia specific clusters. This is because certain genes may not have counterparts in lower ciliary animals but do in higher ciliary species. Visit https://ciliogenics.com to view each gene and heatmaps for clusters. With a combined total of 232 genes, the cilia-specific clusters 31 and 37 contain 159 and 73 putative ciliary candidate genes, respectively. Gene ontology (GO) analysis reveals that Cluster 31 and Cluster 37 have cilia-associated genes. Of these 232 genes, 110 genes are known ciliary genes while the rest are putative ciliary genes, including 36 genes shared by our four human scRNA-seq analyses (**Figure 4E and Table S3**). These 36 genes, including WDR54, ZC2HC1A, and ZNF474, are all likely to be strong ciliary gene candidates because they were found independently by two different approaches. Surprisingly, 311 genes were discovered solely through four human sgRNA-seq analyses, whereas only 86 genes are revealed through comparative genomics analysis, suggesting the independent potential of each method for revealing novel ciliary candidate genes (**Figure S2**).

**Figure 4.**
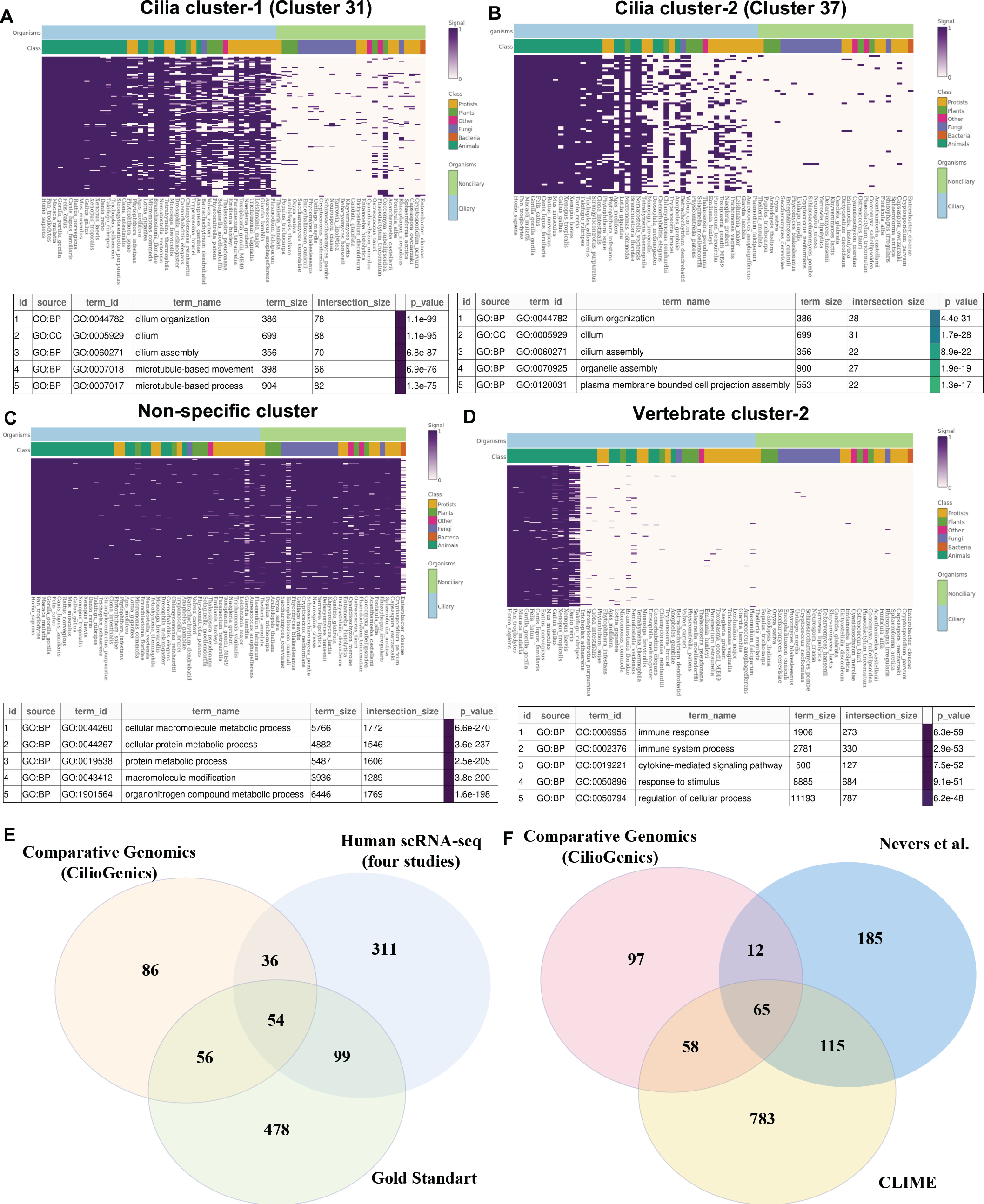
Comparative genomics (phylogenetic profiling) identifies previously unknown ciliary candidate genes. **A, B, C, and D)** A, B, C, and D depict heatmaps of genes in the cilia clusters (31 and 37), non-cilia-specific clusters, and vertebrate-specific clusters. The gene ontology (GO) of each cluster’s genes is provided below the heatmap. **E)** The ciliary candidate genes from CilioGenics (the study) are compared with the ciliary candidate genes from scRNA-seq and the gold standard ciliary genes. Venn diagram displays the findings. **F)** The Venn diagram compares ciliary candidate genes from CilioGenics (this study), CLIME, and Never’s work (Li et al., 2014; Nevers et al., 2017).

### Gene regulatory network and protein-protein interactions (PPI)

Transcription factors and cilia-specific motifs have been extensively used to reveal ciliary candidate genes. Here, we next dig into the transcription factors that are known to regulate them. Chen et al. used scRNA-seq data to identify genetic regulatory networks using nine different species for the lung, and this approach revealed transcription factor (TF)-target interactions (Chen et al., 2021). Here, we have used the transcription factor (TF)-target interactions for ciliated cells, in combination with the scRNA-seq and comparative genomics dataset described above, to identify candidate ciliary genes. Specifically, we chose 6 cilia-related transcription factors (FOXJ1, RFX2, RFX3, JAZF1, GLIS3, and MYB). JAZF1 has emerged as a “regulator of cilia differentiation” and is thought to operate upstream of FOXJ1 (an F-box transcription factor) expression, which is required for constructing motile cilia (Chung et al., 2012; Didon et al., 2013; Johnson et al., 2018; Kang et al., 2009; Tan et al., 2013; Yu et al., 2008). We reasoned that if FOXJ1 functions downstream of JAZF1, the expression of ciliary genes should be influenced by both; thus, we include both in our study (Johnson et al., 2018; Yu et al., 2008). GLIS3 was found to localize to the primary cilium and is required for renal cilium formation, which suggests that it might regulate cilia-related gene expression (Kang et al., 2009).

Our genetic regulatory network analysis (see methods for data analysis) showed GLIS3 is likely involved in the regulation of 562 genes in ciliated cells, which includes 127 Gold Standard ciliary genes. Notably, we used the most recent version of the Gold Standard ciliary gene list for all of our comparisons, and genes that weren’t included in the new Gold Standard list assumed to be unknown (Vasquez et al., 2021). JAZF1 contains 270 cilia-related genes, including 89 Gold Standard ciliary genes, whereas RFX2 has 99 Gold Standard ciliary genes among its 246 genes. RFX3 has 178 Gold Standard ciliary genes out of 878 regulated genes, while MYB has 50 Gold Standard ciliary genes out of 126 genes, and FOXJ1 has 107 Gold Standard ciliary genes out of 433 genes. Taken together, our analysis reveals that 20-40 % of genes regulated by these TF are Gold Standard ciliary genes while the rest are potentially novel ciliary candidate genes (**Figure 5A**). Further analysis revealed that 392 distinct genes are influenced by at least two different ciliary TFs while 6 ciliary TFs affect 16 genes, including ciliary candidate genes (SPAG17, STK33, CCDC170, ERICH3, CCDC146, and CCDC113) and Gold Standard ciliary genes (AK7, ARMC2, ULK4, SYNE1, FANK1, CFAP43, and CCDC39) (**Figure 5B and C, Table S4**). Not surprisingly, CCDC170, ERICH3, CCDC170, CCDC146, and SPAG17 were found to localize to cilia /the base of cilia, suggesting that our TF-based network can uncover novel and known ciliary genes (Merchant et al., 2007; Sigg et al., 2017).

**Figure 5.**
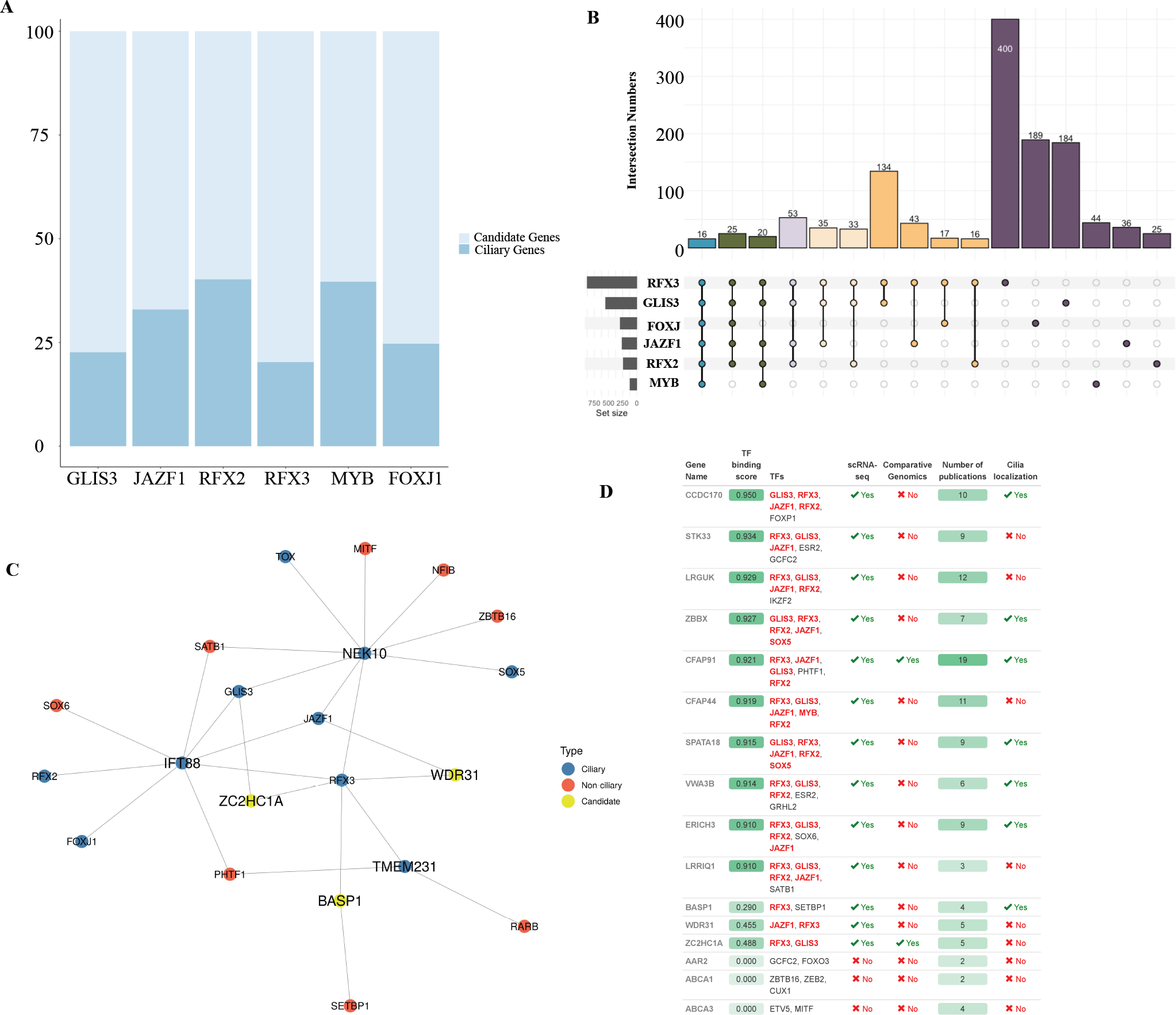
Transcription factor (TF) analysis reveals many new ciliary candidate genes. **A)** The stacked bar graph shows the comparison of FOXJ1, RFX2, RFX3, MYB, GLIS3, and JAZF1 target genes to cilia genes in proportion. Ciliary genes are indicated in dark blue, whereas ciliary candidate genes are labeled in light blue. **B)** The number of intersections of FOXJ1, RFX2, RFX3, MYB, GLIS3, and JAZF1 target genes is shown in the Upset plot. Sharing of TFs, including singles, doubles, triples, quartets, quintuples, and sixes are shown in different colors. **C)** Network analysis shows the binding targets of ciliary TF. Ciliary genes, candidate genes, and non-ciliary genes are shown in blue, red, and yellow, respectively. **D)** The table below lists the top ciliary gene candidates identified using TF analysis. The names of TF linked with genes are shown in red. The table shows which genes have ciliary candidacy based on scRNA-seq and comparative genomics. The table displays the results of a manual search for ciliary localization of ciliary candidate genes.

Next, wedownloaded all publicly available protein-protein interactions (PPI) data from IntAct, BioGRID, and HuRI (Hermjakob, 2004; Luck et al., 2020; Oughtred et al., 2019). We hypothesized that the protein-protein interactions could reveal novel ciliary genes as proteins that function in the same organelle and/or cellular regions like cilia will most likely display intracellular physical interactions (**Figure 6A**). We first categorized the proteins into 3 groups: (i) known ciliary components (ie. in the ciliary Gold Standard), (ii) non-ciliary proteins (ie. in the negative ciliary datasets published by Nevers et al., 2017; SYSCILIA Study Group et al., 2013, Vasquez et al., 2021; and (iii) “unknown” (containing possible ciliary proteins). The size of known ciliary genes (also known as Gold Standard) and negative ciliary genes are 687 and 974, respectively (Nevers et al., 2017; Vasquez et al., 2021). When a candidate protein interacts with a known ciliary protein, it obtains a positive score but does not receive negative scores when it interacts with a negative ciliary protein. We also took into consideration the strength of the protein-protein interaction network for individual candidate proteins in terms of how many known ciliary proteins are present (**Figure 6A**). For example, IFT74 and CC2D2A sub-networks contain many known ciliary proteins (**Figure 6B**). To reduce the number of false negative candidate proteins, we raised the threshold for defining a ciliary protein-protein interaction (PPI) to a value greater than 0.70. Our PPI analysis reveals unique approach 992 proteins, of which 283 (30%) are known ciliary components and < 2% are non-ciliary components (**Figure 6C and Table S5**). CCDC138, CCDC14, and CCDC77 are the top candidate ciliary proteins from the PPI analysis and, indeed, they have already been shown to localize to cilia (Drew et al., 2017) (**Figure 6D**). Focusing on three candidate ciliary proteins s WDR54, ZC2HC1A, and ZNF474 identified in thescRNA and comparative genomics analyses described above, we can see that. ZC2HC1A interacts with known ciliary proteins while the other two proteins do not (**Figure 6E**).

**Figure 6.**
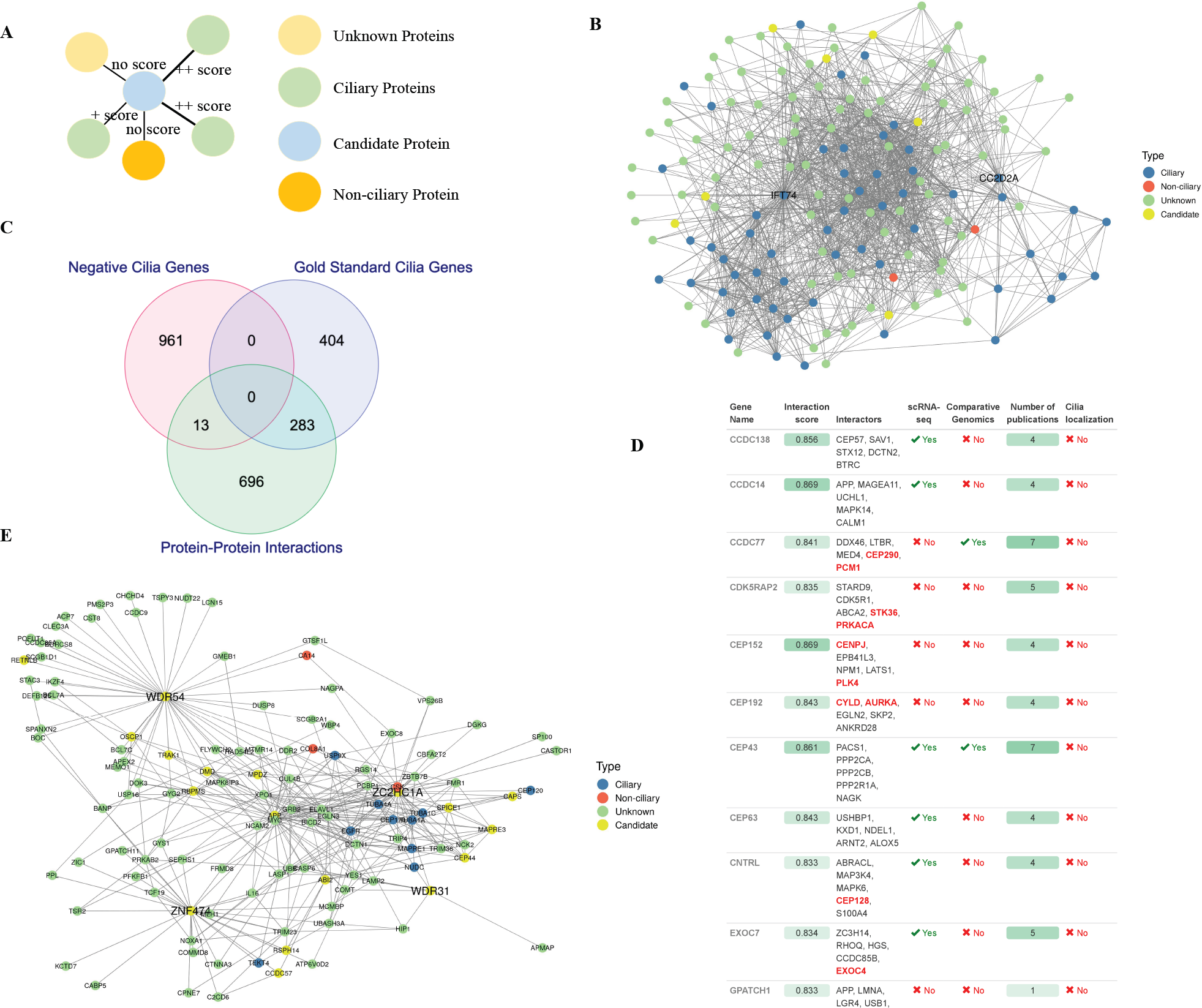
PPIs identify novel ciliary candidate genes. **A)** Shown is the outline of scoring for PPI. Genes are assigned as ciliary, negative, and unknown, all of which are labeled in different colors. Based on its interactions with negative and ciliary genes, each human protein is awarded a negative or positive point. **B)** Network analysis shows PPIs of IFT74 and CC2D2A. Ciliary, negative, unknown, and ciliary candidate genes are represented by the colors red, blue, green, and yellow. **C)** The Venn diagram compares the list of PPI ciliary candidate genes (scores >0.70) to the list of Gold Standard ciliary and negative cilia genes. **D)** PPIs of WDR31, WDR54, ZNF474, and ZC2HC1A are shown in the network. The red, blue, green, and yellow denote ciliary, negative, unknown, and ciliary candidate genes, respectively. **E)** Table displays the top genes from PPIs. The table shows interaction scores, interaction partners, ciliary candidacy from scRNA-seq and comparative genomics, the number of publications found, and cilia localization statutes.

Finally, we downloaded the publicly available Human Protein Atlas (also referred as text mining) database (https://www.proteinatlas.org/) which contains subcellular localization data for most human proteins (Uhlen et al., 2010). Our automated analysis coupled with manual inspection with the Human Protein Atlas showed 370 proteins localize to cilia;75 are “known” ciliary proteins, while the remaining 295 proteins, including BASP1, are poorly characterized novel ciliary (**Table S6**). Taken together, our TF network, protein-protein interaction analysis, and text mining uncover novel ciliary genes.

### Combined method CilioGenics is superior over any single method

Our analysis reveals that the ability of each separate approach to identify known ciliary genes and prospective ciliary candidate genes varies, even though they can all find new ciliary molecular components. Single methods differ in their abilities to identify known ciliary genes and predict ciliary candidate genes (**Figures 7A and B**). TTC39A/C and TMEM145, for instance, have emerged as strong candidate ciliary genes based exclusively on scRNA-seq analysis from *C. elegans*, but not based on scRNA-seq analysis from humans and the other four techniques (PPI interaction, comparative genomics, TF-network analysis, and text mining). We indeed confirm that both TTC39A/C and TMEM145 are localized to the cilia of sensory neurons in the head (amphid) and tail (phasmid) in *C. elegans* (**Figure 7C**). However, this makes choosing a new cilia candidate gene more complicated because of how to prioritize the cilia candidate genes. The combined method—in which all approaches are considered—might be more successful at identifying new cilia genes and weeding out false negative candidates. To obtain more potent results and outcomes that are more biologically instructive, we evaluated the data in the aggregate.

**Figure 7.**
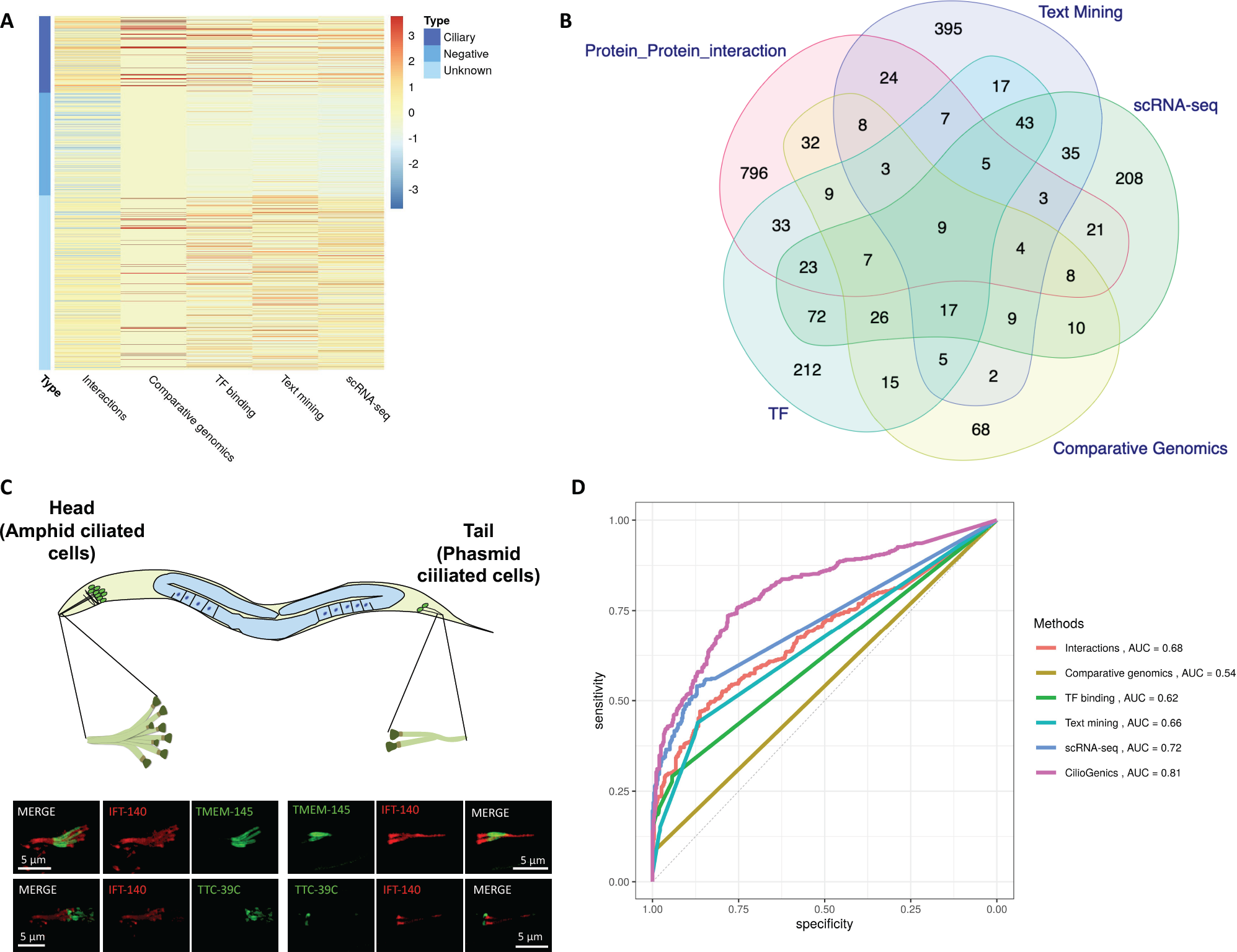
Integrated method CilioGenics is superior to any single method. **A and B)** A heatmap and Venn diagram of ciliary candidate genes from scRNA-seq as well as PPI interaction, comparative genomics, TF-network analysis, and text mining are displayed. No overlap of the putative genes in the heatmap with the negative genes. **C)** Shown are the localization of TMEM-145 and TTC-39c in the head and tails of ciliated sensory neurons (green) in *C. elegans.* IFT-140 (also known as CHE-11, red) is used as the ciliary marker. Scale bar = 5 μm. **D)** Shown is the ROC curve plot displaying the specificity and sensitivity of the single approaches and CilioGenics combined method for predicting ciliary genes. The CilioGenics AUC is superior to any single method.

Initially, the sum scores, called CilioGenics score, are calculated for each human gene’s potential to be a ciliary gene after determining a score for each gene using inputs from scRNA-seq, PPI interaction, comparative genomics, TF-network analysis, and text mining. We next used the AUC-ROC (Area under the Receiver Operating Characteristic Curve) curve to evaluate the performance of our five distinct approaches and CilioGenics. The AUC-ROC displays the sensitivity (true positive rate) on the *y*-axis and specificity (true negative rate) on the *x*-axis. The scRNA-seq analysis has the highest AUC score (0.72), followed by the protein-protein interaction analysis (0.68). Interestingly, the comparative genomics approach has the lowest AUC score (0.54) among all other approaches (**Figure 7D**). The inclusion of the average conservation clusters (1, 4, 6, 9, 16, and 30) into scoring, together with two cilia specific clusters (31 and 37) is most likely the reason for the lowest AUC score for comparative genomics. The AUC-ROC score for CilioGenics is 0.81, suggesting that the combination CilioGenics method is superior to any single method (**Figure 7D**).

### WDR54, ZNF474 and ZC2HC1A are novel ciliary genes

The practical validation of new proposed methods is important to confirm the computationally postulated findings, but the large number of predicted genes limits the extensive experimental execution. To put our CilioGenics method to the test in terms of predicting novel ciliary genes, we focus on the top 500 genes on the CilioGenics gene list. We compared the CilioGenics gene list with the lists of negative genes (975 genes) and Gold Standard ciliary genes (688 genes). Our analysis reveals that the top 500 genes contain 2 negative genes and 242 Gold Standard ciliary genes, suggesting our CilioGenics gene list can successfully identify the Gold Standard ciliary genes. We next focused on these two negative ciliary genes (MUC4 and TUSC3) among the top 500 CilioGenics gene list (**Table S8**). Interestingly, MUC4 has already been discovered to be expressed in ciliated cells and to localize to ciliary shafts (Kesimer et al., 2013). We next concentrate on the other previously unknown 264 ciliary candidate genes in the CilioGenics top 500 gene list (**Figure 8A**). There were already experimental validations for 28 genes, including *BASP1, ANKRD45, DNAH12, IQUB, DYDC2, TEX9, LRRIQ3, BAIAP3, C1orf87, KIF9, EFHC2, MIPEP, C9orf116, ARMC3, PLCH1, C7orf57, EFHB, RIBC2, RGS22, ZBBX, CCDC146, AGBL2, CCDC89, EFCAB1, C21orf58, MDH1B, PPIL6* and *MAP9*, further strengthening the power of our CilioGenics methods in identifying novel ciliary genes (Ivliev et al., 2012) (**Figure 8A**). Notably, none of these genes are used to evaluate CilioGenics’ ability to identify ciliary genes. Furthermore, *WDR54, ZC2HC1A*, and *ZNF474* were among the top 500 gene lists, and we next wanted to carry out subcellular localization analyses in mammals and *C. elegans* to determine whether these non-assigned genes are ciliary. Using this approach, we had already confirmed the cilia localization of WDR31 and ELMOD3 in both human RPE1 cells and *C. elegans*. ZC2HC1A, WDR54, and ZNF474 are listed in the following order on the cilia CilioGenics candidate gene list: 72, 189, and 290, respectively. Confocal microscopy analysis reveals that WDR54, ZC2HC1A, and ZNF474 localize to the cilia of sensory neurons of head and tails in *C. elegans*. Furthermore, endogenous staining with available WDR54 antibody confirms that human WDR54 is indeed in the cilia of human RPE1 cells. Moreover, we selected the 975th-ranked ANKRD26, the *C. elegans* CANK-26 for confirmation. *C. elegans* CANK-26 (the human ANKRD26 ortholog) is enriched at the basal body of cilia (**Figure 8B**).

**Figure 8.**
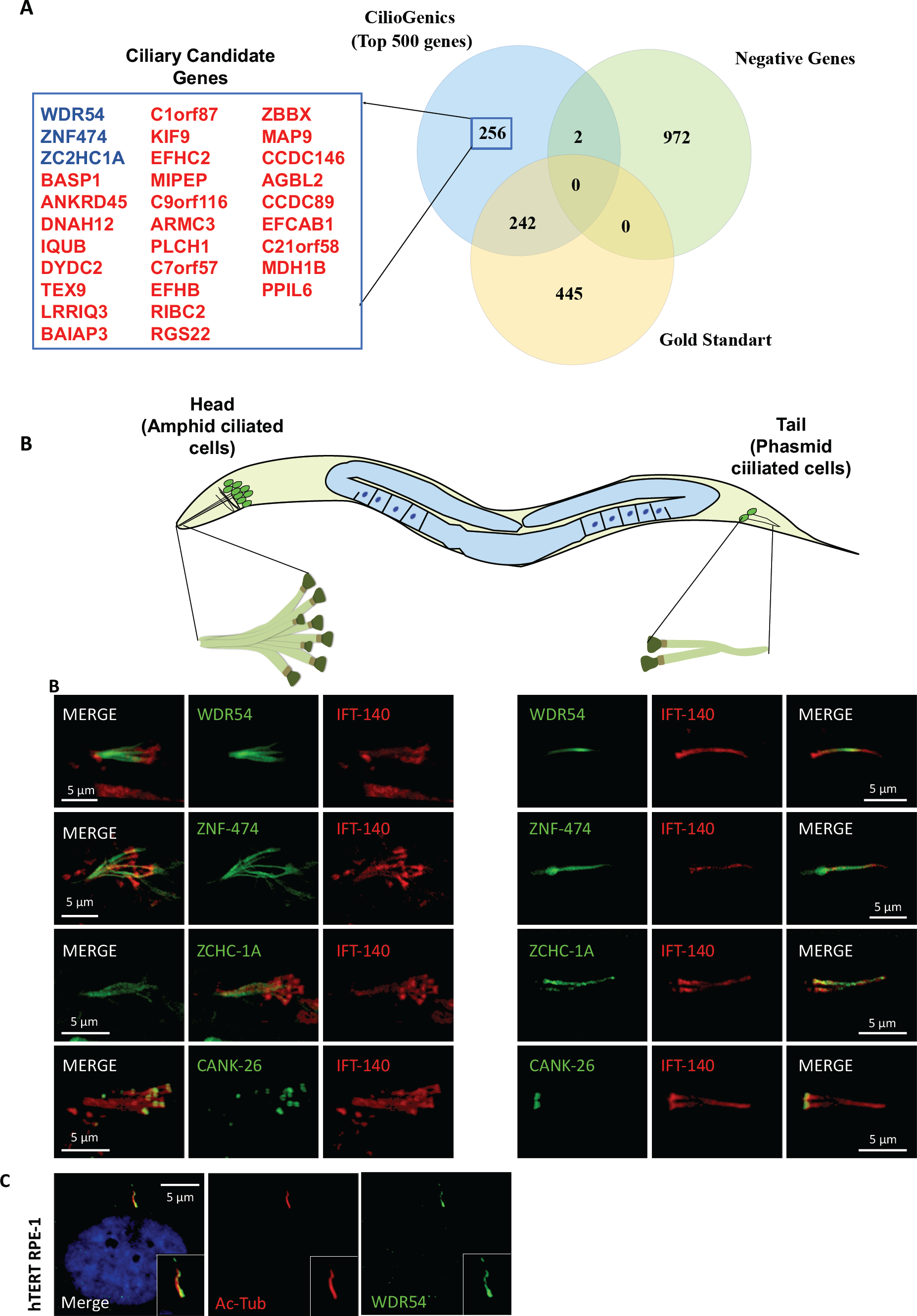
WDR54, ZC2HC1A, and ZNF474 localize to cilia. **A)** The Venn diagram compares the top 500 CilioGenics genes to negative and Gold Standard ciliary genes. The ciliary candidate genes, whose protein products are shown to localize to cilia are shown in red and blue **B)** Fluorescence images display the cilia localization of CANK-26 (the human ANKRD26 ortholog), ZCHC-1A, WDR-54, and ZNF-474 (green) in the head and tails in *C. elegans.* IFT-140 (read) marks cilia. Localization of WDR54 (green) acetylated tubulin (Ac-Tubulin, red), and nucleus (blue) are shown in human retinal pigment epithelial-1 (RPE-1) cells. Scale bar = 5 μm.

## Discussion

More than 50 different papers contributed to the list of ciliary candidate genes, but even with their unquestionable contribution in the discovery of many ciliary genes, the complete list of ciliary genes is far from complete. SYSCILIA Gold Standard (SCGSv2) publication provided the list of 686 known ciliary genes.

Consistent with this foresight, while the 2019 CiliaCarta work suggested there are 956 prospective ciliary genes, a 2021 updated SYSCILIA Gold Standard (SCGSv2) publication provided the list of 686 known ciliary genes (Andersen et al., 2003; Arnaiz et al., 2010; Avidor-Reiss et al., 2004; Baron et al., 2007; Blacque et al., 2005; Boesger et al., 2009; Boldt et al., 2016; Breslow et al., 2018; Broadhead et al., 2006; Cao et al., 2017; Choksi et al., 2014; Efimenko et al., 2005; Fritz-Laylin and Cande, 2010; Geremek et al., 2014, 2011; Hodges et al., 2011; Hoh et al., 2012; Ishikawa et al., 2012; Ivliev et al., 2012; Jakobsen et al., 2011; Keller et al., 2005; Kilburn et al., 2007; Kim et al., 2010; Kubo et al., 2008; Lambacher et al., 2016; Lauwaet et al., 2011; Li et al., 2004; Liu et al., 2007; May et al., 2021; Mayer et al., 2009, 2007; McClintock et al., 2008; Merchant et al., 2007; Mick et al., 2015; Müller et al., 2010; Nakachi et al., 2011; Nogales-Cadenas et al., 2009; Ostrowski et al., 2002; Pazour et al., 2005; Phirke et al., 2011; Reyfman et al., 2019; Roosing et al., 2015; Ross et al., 2007; Sigg et al., 2017; Stubbs et al., 2008; UK10K Consortium et al., 2015; van Dam et al., 2019; Vasquez et al., 2021; Yano et al., 2013). Many of these 57 independent works each used a different approach to find ciliary candidate genes, but their capacity to identify the same set of genes differs from one another (**Table S7**). Comparison of the published lists of ciliary candidate genes reveals the names of the human genes identified by each paper, which allows us to rank genes based on identification of a number of works. 12268 different genes were picked up as ciliary candidate genes by at least one study, whereas 5782 and 2772 genes were chosen as ciliary candidate genes by only one and two studies, respectively. The gene ranking reveals that the first 44 genes are identified as ciliary candidate genes by only 20 publications; the other 37 studies failed to do so. This statistic alone shows how differently each work is able to disclose the ciliary candidate genes, but also proves it has been problematic for all researchers and clinical geneticists to prioritize the ciliary candidate genes whenever they look for the genes that cause disease. Furthermore, the independent gene list produced by our five separate approaches is consistent with the aforementioned conclusion; each has a distinctive ability to discover both known and undiscovered ciliary genes. A strategy incorporating many approaches would likely perform better than each individual approach used alone.

Consistent with expectation, for identifying known ciliary genes and predicting the novel ciliary genes, our novel combined method (CilioGenics) outperforms all other methods evaluated. In the current study, building CilioGenics scores for the ciliary potential of each human gene takes into account five different methods, including (1) protein-protein interactions (PPI) scores obtained by analyzing and merging protein-protein interaction data from IntAct, BioGRID, and HuRI, (2) scores for scRNA-seq data, which analyze fours scRNA-seq data from human lungs and *C. elegans*, (3) scores for comparative genomics, which make a comparison of 72 ciliary and non-ciliary organisms, (4) score from TF-network analysis, which use the binding targets of FOXJ1, RFX2, RFX3, MYB, GLIS3, and JAZF1 in ciliated cells, and (5) text mining scores, which come from the Human Protein Atlas. CiliaCarta integrated multiple datasets to predict the ciliary candidate genes (van Dam et al., 2019). Our primary contribution to CilioGenics compared to the previously available methodologies and CiliaCarta is the integration of more diverse datasets, such as multiple scRNA-seq, comparative genomics, Human Protein Atlas, and different sets of TFs. We demonstrate that the integrated approach is superior for predicting the potential ciliary candidate genes and eliminating false positive candidate genes.

Our comparative genomics analysis reveals that *WDR54*, *ZNF474*, and *ZC2HC1A* show up as ciliary candidate genes, but none of the previous comparative genomics analyses suggested them as ciliary candidate genes. The failure of other studies to recognize them as ciliary gene candidates may have been partially explained by organism choices, use of different thresholds, and analysis types (Avidor-Reiss et al., 2004; Li et al., 2014; Nevers et al., 2017). Nonetheless, these three genes showed up in our scRNA-seq study as ciliary candidate genes, which increases our confidence that they are ciliary genes. Our combined CilioGenics technique placed them in the top ciliary candidate genes, therefore we gave them precedence and verified their cilia localizations.

Can CilioGenics accurately predict the total number of ciliary genes? This would be a challenging question to answer and depends on where the threshold will be cut. Furthermore, the current version of CilioGenics uses scRNA-seq data from human lung tissue for the prediction, and ciliary genes that are not expressed in the human lung will be apparently missed. We believe that more diverse tissue of scRNA-seq data is needed to increase the range of prediction from the scRNA-seq data. Protein-protein interactions of many human proteins are poorly characterized, and they are not well represented in the CilioGenics scoring. In addition, many of the antibodies utilized for subcellular localization studies of poorly characterized human proteins were far from ideal, and future improvement in antibody technology will be helpful in this area. Nonetheless, we are confident that the number of genes in human cilia is greater than 687, which is the number that is presently suggested as the Gold standard ciliary genes. This idea is supported by the fact that 295 previously uncharacterized proteins from the Human Protein Atlas have localizations specific to cilia, bringing the total number of ciliary genes to 982. Furthermore, the current study also validates the localisation of 5 additional ciliary genes. In summary, we believe that the CilioGenics in conjunction with our up to date CilioGenics website (https://ciliogenics.com/) would be helpful to scientists to search for ciliary candidate genes, and they can use the provided dataset to prioritize their candidate genes. Additionally, users can find the number of publications where a gene was proposed as a ciliary candidate gene, check presence of gene in different types of comparative genomics clusters (cilia organisms specific clusters, average conservation cluster, or low specify cluster), or expression patterns of genes in the human lungs. The current version of CilioGenics and the website will be openly updated and other data sets, including scRNA-seq of other tissues will be integrated.

## Materials and Methods

### Human orthologs of *C. elegans* genes

As a part of our previous work, we had generated the *C. elegans* orthologs of human genes (Pir et al., 2022; The Alliance of Genome Resources Consortium et al., 2020).

### scRNA-Seq analysis of human Lungs and single-cell RNA sequence analysis of *C. elegans*

The human lung scRNA-seq raw data were retrieved from the Gene Expression Omnibus (www.ncbi.nlm.nih.gov/geo). The following accession number was used to download the raw data: (GSE122960) (Reyfman et al., 2019), followed by analysis with the Seurat R package (Stuart et al., 2019). Default settings were used and codes for analysis can be found at https://github.com/thekaplanlab/CilioGenics_Analysis. The three pre-analyzed data for the human lung scRNA-seq were obtained as .RDS file (Carraro et al., 2021; Habermann et al., 2020; Kadur Lakshminarasimha Murthy et al., 2022). For *C.* elegans scRNA-seq, the RDS file for the pre-analysed scRNA-seq data was downloaded (Cao et al., 2017). Default settings were used to conduct differential expression analysis between ciliated and other cellular clusters. **Supplementary Table 1 and Table 2** contain ciliary candidate gene names from *C. elegans* scRNA-seq and Human scRNA-seq.

### BLAST analysis for 72 organisms and Clustering

The list of ciliary and non-ciliary organisms were chosen and their genomes were downloaded from NCBI using our in-house codes. The list of organisms with download links were provided in the **Supplementary Table 11**. The longest protein transcript for each gene was selected, followed by performing Basic local alignment search tool (BLAST) (Ye et al., 2006), and searching the orthologs of each human gene among 70 different organisms. The emerging file was filtered by selecting a single transcript for each gene with the highest bit scores. As a final filter, genes with p-values lower than 0.001 and query cover higher than 50% were chosen. Finally, if human genes have orthologs in the other organisms, they received a score of 1, otherwise they received a value of 0. The final file was used for clustering. In a brief, a dissimilarity matrix was generated using the ‘daisy’ package with the gower metric. The dissimilarity matrix was then subjected to hierarchical clustering in order to produce a tree. The generated trees were partitioned into 40 clusters using the ‘cutree’ function in order to efficiently cluster the data. The genes that have orthologous genes in just ciliated species are found in only two groupings. For each cluster (a total of 40 clusters) we create a heatmap using the pheatmap package. **Supplementary Table 3** contains the list of genes Cluster 31 and Cluster 37. The CilioGenics website provides users with access to names of genes in each cluster (https://ciliogenics.com/).

### Protein-Protein interactions (PPI) Analysis

The protein-protein interaction (PPI) data for IntAct, BioGRID and HuRI were downloaded from the websites corresponding to the data (https://www.ebi.ac.uk/intact/), (https://thebiogrid.org/), and (http://www.interactome-atlas.org/) (Hermjakob, 2004; Luck et al., 2020; Oughtred et al., 2019). For the IntAct, the data without no PPI score (miscore) are excluded. PPI interactions data from organisms, including *Homo sapiens, Mus musculus, Drosophila melanogaster* and *Caenorhabditis elegans* are included for BioGRID and IntAct. For HuRI, *H. sapiens*-related data, on the other hand, were utilized exclusively. Every identifier, including gene identifiers and UniProt identifiers, is converted to an HGNC gene name. Finally, using Alliance of Genome Resources, gene names from non-human organisms are changed to their human equivalents (The Alliance of Genome Resources Consortium et al., 2020). If both the interactors and the publication identifiers are the same, an interaction was considered redundant, and the redundant interactions were when combining all three data. Human gene list coming from PPI can be found at the **Supplementary Table 5**. We used a modified version of MIscore to score interactions, which assigns a score based on the weights of the detection method, the kind of interaction, and the quantity of publications (Villaveces et al., 2015). However, because doing so would only favor the established ciliary genes, we did not include publications when rating.

### Transcription Factor (TF) target interaction prediction

TF-target interaction prediction data were taken from the recent paper, which presented a single cell atlas for 11 different organisms, including rabbit, duck, cat, dog, hamster, lizard, goat, pigeon, pangolin, tiger, and deer (Chen et al., 2021). The predictions based on ciliated cells were taken into account. Six cilia-related transcription factors (TF), including RFX2, RFX3, MYB, GLIS3, and JAZF1, were chosen, and the genes that were expected to interact with these ciliary TFs were anticipated to be ciliary genes (Chung et al., 2012; Didon et al., 2013; Johnson et al., 2018; Kang et al., 2009; Tan et al., 2013; Yu et al., 2008). Furthermore, the predicted FOXJ1 regulatory network gene list was taken from Mukherjee et al. and combined with the other list (Mukherjee et al., 2019). The gene names coming from TF analysis are listed in the **Supplementary Table 4.**

### Text Mining (Protein Atlas)

The website for the Protein Atlas was scanned, and genes with at least one of the keywords “cilia, cilium, flagella, flagellum, and/or centrosome” were collected (Uhlen et al., 2010). Also, those genes were thought to be localized to cilia if at least one of the “positivity or stain” terms appeared alongside cilia or cilium. Otherwise, it was assumed to only be expressed in ciliated cells. The **Supplementary Table 6** contains a list of all gene names discovered using text mining.

### Gene ontology (GO)

The R package gprofiler2 was used to generate gene ontologies for the genes identified in comparative genomics clusters. The relevant figures: Figure 4A, B, C and D (Kolberg et al., 2020).

### Ciliary Gold Standard Genes and Negative Ciliary Genes

Nevers and colleagues presented the negative ciliary genes, whereas the updated ciliary Gold Standard Ciliary Genes were used (Nevers et al., 2017; Vasquez et al., 2021). The **Supplementary Table 9 and 10** contain the gene names of Gold Standard ciliary genes and negative ciliary genes, respectively.

### Collection of Publications

The names, types of organisms, web links, and other information about the papers and tables that contained the list of ciliary candidate genes were gathered from all cilia-related publications and reported in **Supplemental Table 7.**

### Creation of CilioGenics Database

For the comprehensive cilia database, R Shiny was used to develop a database called CilioGenics. All codes and datasets are available on the Kaplan Lab Github: https://github.com/thekaplanlab/CilioGenics-website.

### Strains Maintenance

The *C. elegans* wild type isolate strains (N2) were raised on the previously described Nematode Growth Medium (NGM) (Brenner, 1974).

### Generation of Translational and Transcriptional Transgenic strains

The pPD95.67 plasmid, a modified *C. elegans* expression vector, was used to clone the 500 base pairs (bp) of the gene promoter followed by 1542 bp of the cDNA for C15A7.2 (tmem-145) directly in front of the GFP. The length of a gene’s promoter was limited to 500 base pairs, however the length of each cDNA was as follows: 396 base pairs for cDNA of M153.4 (znf-474), 1044 base pairs for cDNA of F39H12.2 (wdr-54), and 969 base pairs for cDNA of T03G11.3 (zchc-1a). SphI and AgeI restriction enzymes were used in the cloning. The resulting plasmids were as follow: wdr-54promoter::WDR-54::GFP; tmem-145promoter::TMEM-145::GFP, znf-474promoter::ZNF-474::GFP, and zchc-1a::ZCHCC-1A::GFP. With the exception of tmem-145, All of the other genes and 1380 bp of cDNA of K10G6.4 (cank-26) were cloned between 500 base pairs long *arl-13* promoter (a cilia specific promoter) and GFP. The following plasmids were produced:: arl-13promoter::WDR-54::GFP; arl-13promoter::ZNF-474::GFP, arl-13::CANK-26::GFP, and arl-13a::ZCHC-1A::GFP. The TTC-39c::GFP (C32D5.6) fosmid were purchased from Source BioScience. Unless stated, a microinjection of 50 ng/μl RF4 (Roller selection marker) at a dosage of 5-25 ng/μl was performed for each construct. The resulting transgenic strains were crossed into Intraflagellar transport 140 (IFT-140)::mCherry (known as CHE-11), an IFT-A component used as a ciliary marker.

### Score Calculation and ROC curve (receiver operating characteristic curve)

Five independent approaches, including scRNA-seq, PPI interaction, comparative genomics, TF-network analysis, and text mining generated a list of ciliary candidate genes. The ciliary capacity of each gene was determined by calculating scores from each approach. In scoring of the PPI interaction, the nature of interactions, such as direct interaction, biochemical, protein complementation assay, post transcriptional interference, genetic interaction, physical association were considered. Biogrid, intact and HuRi were analyzed, and duplicated interactions were removed. Here are scoring: MI:0013: biophysical: 1, MI:0090: protein complementation assay: 0,66, MI:0254: genetic interference: 0.10, MI:0255: post transcriptional interference: 0.10, MI:0401: biochemical: 1, MI:0428: imaging technique: 0.33, MI:0208: genetic interaction (sensu unexpected): 0.1, MI:0403: colocalization: 0.33, MI:0914: association: 0.33, MI:0915: physical association: 0.66, MI:0407: direct interaction: 1.

The TF-network scoring: The six transcription factors (TF) RFX2, RFX3, MYB, GLIS3, and JAZF1, FOXJ1 were chosen, and points were assigned to any gene that was a target of these TF, followed by normalization.

scRNA-seq scoring: Cilia-specific expressing genes from human lung scRNA-seq (Reyfman, Carraro, and Habermann) and *C. elegans* scRNA-seq studies were scored and normalized.

Comparative genomics scoring: Clustering results of comparative genomics of 72 different organisms leads to 40 different clusters. Visual inspection leads to identification of two cilia specific clusters (31 and 37), average conservation clusters (1, 4, 6, 9, 16, and 30) and low specificity clusters (2, 3, 5, 7, 8, 10, 11, 12, 13, 14, 15, 17, 18, 19, 20, 21, 22, 23, 24, 25, 26, 27, 28, 29, 31, 32, 33, 34, 35, 36, 37, 38, 39, and 40). Genes in the two cilia-specific clusters are assigned a score of 1, while genes in the average conservation and low specificity clusters are assigned 0.5 and 0 points, respectively.

Text mining scoring: Genes that encode cilia localizing proteins receive a score of 1, whereas genes encoding cilia expressing proteins receive a value of 0.5. Genes that are retrieved with “centrosome” receive 0.25.

The accuracy of each method was determined using a randomly generated test and trained gene lists, leading to the generation of a confusion matrix. The confusion matrix (cilia vs non-cilia) was used to calculate F1 scores for each approach.

Precision = *TP* / (*TP* + *FP*)

Recall: = *TP* / (*TP* + *FN*)

F1 Score = 2

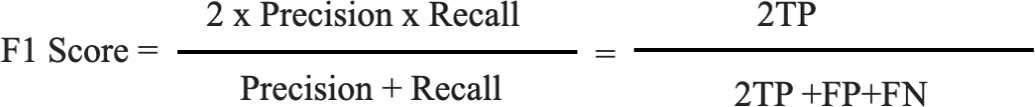

Number of true positives (TP)

Number of false positives (FP)

Number of false negatives (FN)

F1 scores were determined for each method. With help of F1 scores, the CilioGenics scores (sum scores) for each gene were calculated. The accuracy of calculated scores was determined by ROC-AUC curve generation for each method, including CilioGenics.

### Cell Culture

hTERT RPE-1 (ATCC-CRL-4000) cells were were maintained in DMEM:F-12 media (Gibco, 11320033) supplemented with 10% FBS (Gibco, 26140079) and 1% Pen-Strep (Gibco, 15070063) at 37 °C in 5% CO_2_ (Oner et al., 2013).

### Immunostaining

Cell Imaging— hTERT RPE-1 cells were seeded onto sterile 24-mm polylysine-D-coated coverslips in 6-well plate, maintained in 10% FBS for 24 hours, later switched to serum free medium for 48 hours to induce ciliogenesis.

Serum starved hTERT RPE-1 cells on poly-D-lysine-coated glass coverslips were washed twice with cell-washing solution (CWS; 2.6 mM KCl, 137 mM NaCl_2_, 10 mM Na_2_HPO_4_, 1.8 mM KH_2_PO_4_). Then cells were fixed with 4% paraformaldehyde - 4% sucrose in CWS for 10 min. Each well was washed three times with CWS and permeabilized by 5min incubation with 0.2% Triton X-100 in CWS. Each coverslip was blocked for 60 min. incubation with 10% normal goat serum (İnvitrogen, 50197Z) in CWS. Cells were incubated with Anti-WDR-54 antibody (1/100, Invitrogen, PA5-62806) and anti-acetylated tubulin (1/150, MilliporeSigma, MABT868) in 1%BSA containing CWS. Goat anti-rabbit AlexaFluor488 and anti-mouse Alexa-Fluor594 (Invitrogen, A32731, A32742) secondary antibodies were diluted by 1/500 into 1% BSA containing cell washing solution. All antibody dilutions were centrifuged at 10,000Xg for 10 min prior to use. The nucleus was stained with 1 mg/ml DAPI at the last washing step for 5 min. Slides were then mounted with glass coverslips by using anti-fade reagent (Invitrogen, P36930) (Vural et al., 2010).

### Coding and Files

The codes used for data analysis and constructing CilioGenics website may be accessed at the respective GitHubs: https://github.com/thekaplanlab/CilioGenics_Analysis.

## Supplementary Files

**Supplementary Figure 1**

Shown is a drawing of the human lung, with cells from trachea/bronchi, bronchioles to alveoli and alveoli. Clara, goblet and multiciliated epithelial cells are shown.

**Supplementary Figure 2**

The Venn figure compares gene lists from four scRNA-seq studies (Carraro, Reyfman, Habermann, and Murthy), comparative genomics (Cluster 31 and Cluster 37), negative, and gold standards.

**Supplementary Figure 3 and Supplementary Figure 4**

Networks comprising the top 500 genes from CilioGenics (**Figure S3**) and 687 genes from Gold Standard (**Figure S4**) are shown. The red, blue, green, and yellow denote ciliary, negative, unknown, and ciliary candidate genes, respectively.

**Supplementary Figure 1**

UMAPs from human lung scRNA-seq are shown after a re-analysis of scRNA-seq (Kadur Lakshminarasimha Murthy et al., 2022).

**Supplementary Table 1:**

The list of ciliary candidate genes from *C. elegans* scRNA-seq

**Supplementary Table 2:**

The human scRNA-seq list of ciliary candidate genes from Carraro, Reyfman, Habermann, and Murthy. Indeed, ciliary genes are indicated. Carraro, Reyfman, Habermann, and Murthy are compared and shown in the gene list. The number of shares and the names of the studies were provided.

**Supplementary Table 3:**

The list of ciliary candidate genes (only cluster 31 and 37) from comparative genomics. Known ciliary genes and ciliary candidate genes from scRNA-seq are labeled.

**Supplementary Table 4:**

List of gene targets of six cilia-related transcription factors (TF), including RFX2, RFX3, MYB, GLIS3, and JAZF1 and FOXJ1, is shown. The known ciliary genes are marked.

**Supplementary Table 5:**

Top candidate genes from the protein-protein interaction (PPI) data for IntAct, BioGRID, and HuRI (score >0.7). PPI’s top candidate genes are compared to a list of negative and gold standards. All genes (11229 genes) from PPI’s are provided..

**Supplementary Table 6:**

List of genes from text mining from Protein Atlas. The known ciliary genes are labeled yes.

**Supplementary Table 7:**

The list includes the names of the genes as well as the number of articles that have identified them as ciliary candidate genes. The publication names are shown. Documentation on chosen articles are supplied, including publication year, file names, organism kinds, and paper webpage.

**Supplementary Table 8:**

Each gene receives a score from each of the five approaches we presented, and the sum of the scores for each gene was determined. Genes are ordered depending on the total of their scores. Top 500 genes from CilioGenics are presented in **Table S8**.

**Supplementary Table 9:**

Gold standard gene list was acquired from Vasquez et al (Vasquez et al., 2021).

**Supplementary Table 10:**

Negative gene list was obtained from Nevers et al (Nevers et al., 2017)

**Supplementary Table 11:**

The organism names used in comparative genomics are listed. Additionally, the protein sequences for each organism are retrieved from NCBI, and download links are provided.

**Supplementary Table 12:**

The list of strains used in the paper is provided.

## Author contributions

MSP handled the majority of the bioinformatic work, whereas FY and SC generated transgenic strains and performed microscopy analysis. SSO performed immunostaining in RPE cells. ENFK and OEB contributed reagents. OIK directed the project, produced the figures, and wrote the manuscript with input from all authors.

## Acknowledgments

We thank the Kaplan Lab members for their valuable inputs.

